# Quantification of microbubble-induced streaming across ultrasound frequencies using particle image velocimetry

**DOI:** 10.64898/2026.01.07.698213

**Authors:** Reza Rasouli, Brad Hartl, Soren Konecky

## Abstract

Functional microbubbles are widely used in therapeutic ultrasound for drug delivery, blood-brain barrier opening, clot lysis, and other biomedical interventions. Their therapeutic effects arise largely from acoustic microstreaming, which generates localized shear stresses and enhances transport around oscillating bubbles. Despite the central role of microstreaming, its dependence on ultrasound excitation conditions is not well defined. Existing studies use diverse and often incomparable frequencies, pressures, and duty cycles, leaving a gap in systematic understanding of how these parameters govern the streaming velocity and shear fields produced by functional microbubbles under physiologically relevant conditions. In this study, we employ particle image velocimetry (PIV) in a popliteal-vein lab-on-chip model to directly quantify the microstreaming velocity and shear-rate fields generated by microbubbles across a wide range of ultrasound conditions including frequency, duty cycle, and acoustic pressure. Measurements at 150 kHz, 300 kHz, 500 kHz, and 1 MHz reveal a pronounced frequency dependence. Counterintuitively, the strongest microstreaming is observed at 150 kHz, with progressively weaker streaming at 300 and 500 kHz, and the weakest flows at 1 MHz. This trend contrasts with the common assumption that microbubble oscillations peak near their ∼1 MHz resonance frequency. Duty cycle further modulated the response: 10% duty cycle generated the weakest fields, whereas 100% duty cycle produced maximal streaming. The results reveal a strong nonlinear pressure dependence at low frequencies, particularly at 300 kPa where flow amplification was most pronounced. These findings underscore that acoustic frequency plays a substantial role in intensity of microbubble-induced streaming, with low-frequency excitation in the 150-300 kHz range producing markedly more efficient streaming and associated shear. Adopting acoustic parameters that maximize microstreaming within this low-frequency window while avoiding conditions that promote inertial cavitation may improve both the efficacy and safety of microbubble-mediated treatments.

## Introduction

Ultrasound-driven microbubbles are powerful agents in biomedical engineering, known for their ability to enhance processes such as clot dissolution, localized drug delivery, and blood brain barrier disruption. ^1–3^ When exposed to an acoustic field, microbubbles undergo volumetric oscillations that substantially amplify the mechanical effects of ultrasound on surrounding tissues and fluids. ^4–6^ A key mechanism by which microbubbles impose therapeutic effects is acoustic microstreaming, which is the steady circulatory flow generated in the fluid around an oscillating bubble. These micro-vortices create localized shear stresses and convective transport that enhance mass transfer, disrupt structural barriers, and increase penetration of therapeutic agents into targets such as thrombi or tissue^5,7,8^. Unlike the extreme, transient flows produced by inertial cavitation (bubble collapse), microstreaming arises from stable bubble oscillations. This phenomenon, also known as stable cavitation, underlies a wide range of therapeutic applications and can be effectively harnessed when acoustic parameters are carefully fine-tuned to sustain and control it. For instance, stable oscillatory microbubbles can reversibly permeabilize cell membranes through microstreaming-generated shear flows, facilitating drug delivery without necessarily causing large-scale bubble collapse^4,9,10^. In clot dissolution, microstreaming around oscillating bubbles increases penetration of lytic enzymes into the thrombus and exposes new binding sites, thereby accelerating fibrinolysis.^3,5,11^

Although, microbubble-mediate acoustic streaming has been studied across a range of biomedical applications, its dependence on ultrasound frequency remains inadequately characterized. The literature employs widely varying frequencies, pressures, and duty cycles, resulting in inconsistent reports of microbubble performance. The resonance frequency of commercially available contrast microbubbles (110 μm diameter) lies in the low MHz range. Indeed, many therapeutic ultrasound protocols historically employed frequencies around 1 MHz under the assumption that near-resonant driving produces the largest bubble oscillations and thus the greatest effect. ^4,8,12^ Paradoxically, some recent findings challenge this assumption. Ilovitsh et al.^13^ observed that standard lipid-shelled microbubbles subjected to a low-frequency 250 kHz ultrasound exhibited greater expansion than at 1 MHz, with expansion ratios reaching ∼30-fold at 250 kHz versus only ∼1.6-fold at 1 MHz under comparable conditions. This suggests that maximum bubble response and by extension, the strongest microstreaming may occur at frequencies well below the linear resonance. Similarly, reports from sub-MHz (∼220-260 kHz) bloodbrain barrier opening studies likewise highlight that lower frequencies may more effectively excite microbubbles and facilitate stable cavitation while reducing attenuation and minimizing destructive collapse. ^14–17^

Another important consideration is the distinction between bubble oscillation amplitude and therapeutic efficacy. While bubble vibration amplitude reflects cavitation activity, it does not directly quantify the streaming-induced shear and mixing, which in many cases is more closely linked to functional outcomes such as drug uptake or thrombolysis. Critically, oscillation amplitude by itself is not an adequate predictor of microstreaming because microstreaming also depends on the angular frequency of excitation and the thickness of the viscous boundary layer, both of which determine how effectively oscillatory motion is converted into steady streaming. Thus, it is crucial to characterize microstreaming itself.

Here we address this gap by directly quantifying interaction of oscillating microbubbles with the surrounding fluid using particle image velocimetry (PIV). PIV offers a direct way to visualize and quantify streaming flows around bubbles by providing spatially resolved measurements of the streaming velocity and shear fields. Unlike indirect approaches such as passive cavitation detection or acoustic backscatter analysis which infer bubble behavior from emitted acoustic signals, PIV captures the actual fluid motion generated by the bubble, enabling precise characterization of the mechanical environment responsible for drug transport, clot disruption, and other therapeutic effects. High-speed optical imaging can visualize bubble oscillations, but it does not quantify the resulting flow, while numerical simulations rely on assumptions about bubble dynamics and fluid properties. In contrast, PIV directly measures the steady flows associated with stable cavitation, offering a robust and unbiased method for assessing how acoustic driving conditions translate into functional microstreaming.^18–22^

In this manuscript, we systematically investigate how ultrasound frequency, duty cycle, and acoustic pressure influence microbubble-induced streaming in a physiologically relevant vein lab-on-chip model. To comprehensively span the parameter space typically used in biomedical ultrasound, we examine four frequencies (150 kHz, 300 kHz, 500 kHz, and 1 MHz), three duty cycles (10%, 20%, and 100%), and three acoustic pressures (100, 200, and 300 kPa). These values encompass the range commonly reported in therapeutic studies while capturing the sub-MHz regime where stable cavitation and strong bubble-fluid interactions have been observed. Through this study, we aim to establish a quantitative basis for selecting ultrasound frequencies and excitation conditions that most effectively harness microbubble-induced streaming for biomedical applications.

## Materials and Methods

### Lab-on-a-Chip (LOC) Device Fabrication

The experimental platform was built using a custom-fabricated lab-on-a-chip (LOC) device designed to simulate a 6 mm diameter popliteal vein, chosen for its clinical relevance in transcutaneous ultrasound targeting. The molds were created using Onshape CAD software (PTC Inc., USA) and manufactured via digital light processing (DLP) 3D printing. Channels were cast in Momentive RTV615 silicone elastomer and bonded to a 20 μm silicone sheet (Gteek, UK) using plasma surface activation.

### Ultrasound Exposure Setup

Ultrasound experiments were performed under static conditions inside a custom-designed acoustic water bath housing the vein-on-chip (LOC) device, the focused ultrasound transducer, and the mounting assembly (Figure 1). The transducer was positioned at the base of the bath, and the LOC device was secured in a 3D-printed holder that aligned the vein-mimicking test chamber with the acoustic focal plane. Degassed water served as the acoustic coupling medium.

**Figure 1.**
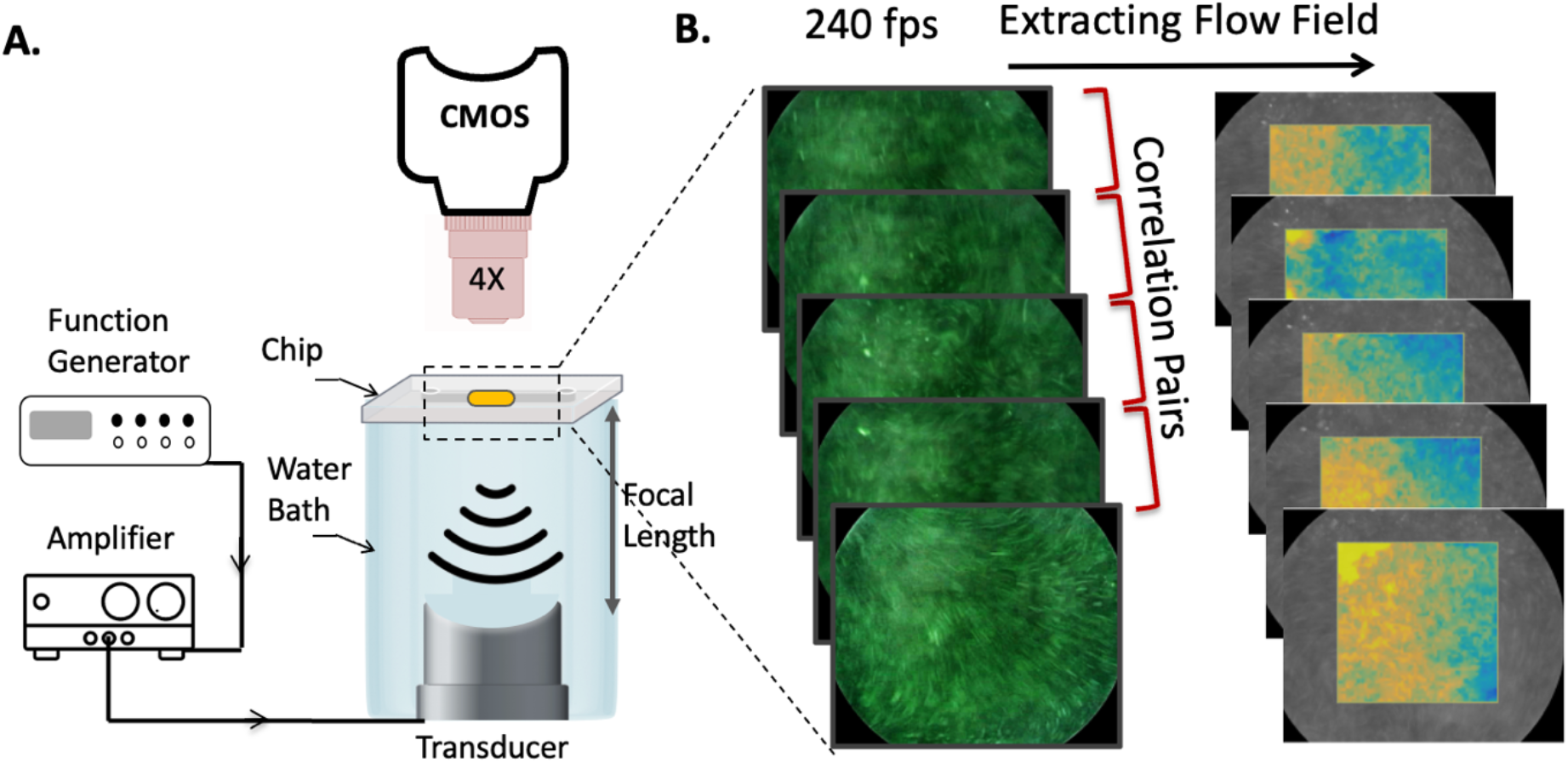
**(A)** Schematic of experiment setup showing the focused ultrasound transducer, acoustic water bath, and inverted microscope used for imaging. The LOC chamber was seeded with 15 μm tracer particles for PIV measurements. **(B)** PIV analysis workflow illustrating sequential image acquisition from the LOC chamber, cross-correlation of image pairs, and extraction of velocity and shear-rate fields used to quantify microbubble-driven microstreaming.

Ultrasound excitation was applied at four frequencies: 150 kHz, 300 kHz, 500 kHz, and 1 MHz using continuous-wave signals with duty cycles of 10%, 20%, or 100% and acoustic pressures of 100, 200, or 300 kPa. A function generator (DG4162, Rigol Technologies) provided the driving waveform, and an RF amplifier (1040L, Electronics & Innovation) controlled the output voltage.

A focused transducer (Benthowave Instrument Inc., Model BII-7652; 60-mm aperture, 36-mm focal length) produced a focal beam that fully covered the 6-mm LOC chamber at all tested frequencies, ensuring uniform insonation across the sample volume.

Vevo MicroMarker microbubbles (FUJIFILM VisualSonics) were reconstituted according to manufacturer instructions and diluted **1:500** in phosphate-buffered saline (PBS). The PBS was selected due to its osmolarity and compatibility with phospholipid-shelled microbubbles. As previously reported, fluid properties such as gas content and ionic strength can modulate microbubble behavior under ultrasound.^23^

### PIV Acquisition and Processing

Microstreaming flow fields were quantified using particle image velocimetry (PIV). The working fluid was seeded with fluorescent polystyrene microspheres (MagSphere Inc., USA) with a nominal diameter of 15 μm, providing sufficient light scattering for high-contrast imaging while remaining small enough to faithfully follow the steady streaming flows with minimal inertial lag.

Experiments were performed on an inverted Revolve microscope (Echo Laboratories, USA) equipped with an iPad-based imaging system. PIV image sequences were acquired in bright-field mode at 240 frames per second to acquire adequate temporal resolution and capture the evolution of the microstreaming patterns generated under ultrasound excitation.

Recorded image sequences were exported and analyzed in PIVlab (MATLAB), which utilizes a multi-pass FFT-based cross-correlation approach. Prior to correlation, images were pre-processed using standard PIVlab operations, including contrast enhancement (CLAHE) to improve particle visibility and reduce illumination nonuniformities. Velocity fields were computed using a three-pass interrogation scheme (128 × 128 px, 64 × 64 px, and 32 × 32 px with 50% overlap) to allow progressive refinement of spatial resolution while maintaining robust correlation in the initial passes. The resulting displacement fields were converted to velocity fields by dividing by the calibrated pixel size and the inter-frame time interval to get two-dimensional velocity maps for each frame.^18,24^

Shear rate was computed directly from the velocity-gradient tensor using PIVlab’s internal formulation. PIVlab reports the symmetric shear component of the tensor, defined as:

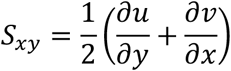

where *u* and *v* are the in-plane velocity components. ^18,24^ To capture peak shear magnitudes while minimizing sensitivity to noise or isolated vector errors, the 95th-percentile absolute shear (P95Shear) within the defined region of interest was extracted for each acoustic condition and used as the primary metric for statistical analysis.

## Results

To characterize how ultrasound driving conditions modulate microbubble-induced microstreaming, we quantified both the velocity fields and shear-rate fields across all tested frequencies, duty cycles, and acoustic pressures.

### Velocity field of the streaming flow

Figure 2 shows representative velocity-magnitude maps at 300 kPa, overlaid on the bright-field image of the vein-on-chip chamber, for four ultrasound frequencies (150 kHz, 300 kHz, 500 kHz, and 1 MHz) and three duty cycles (10%, 20%, and continuous-wave, CW). The grey background corresponds to the imaged chamber region, while the central square indicates the PIV interrogation window, with color denoting the local velocity magnitude.

**Figure 2.**
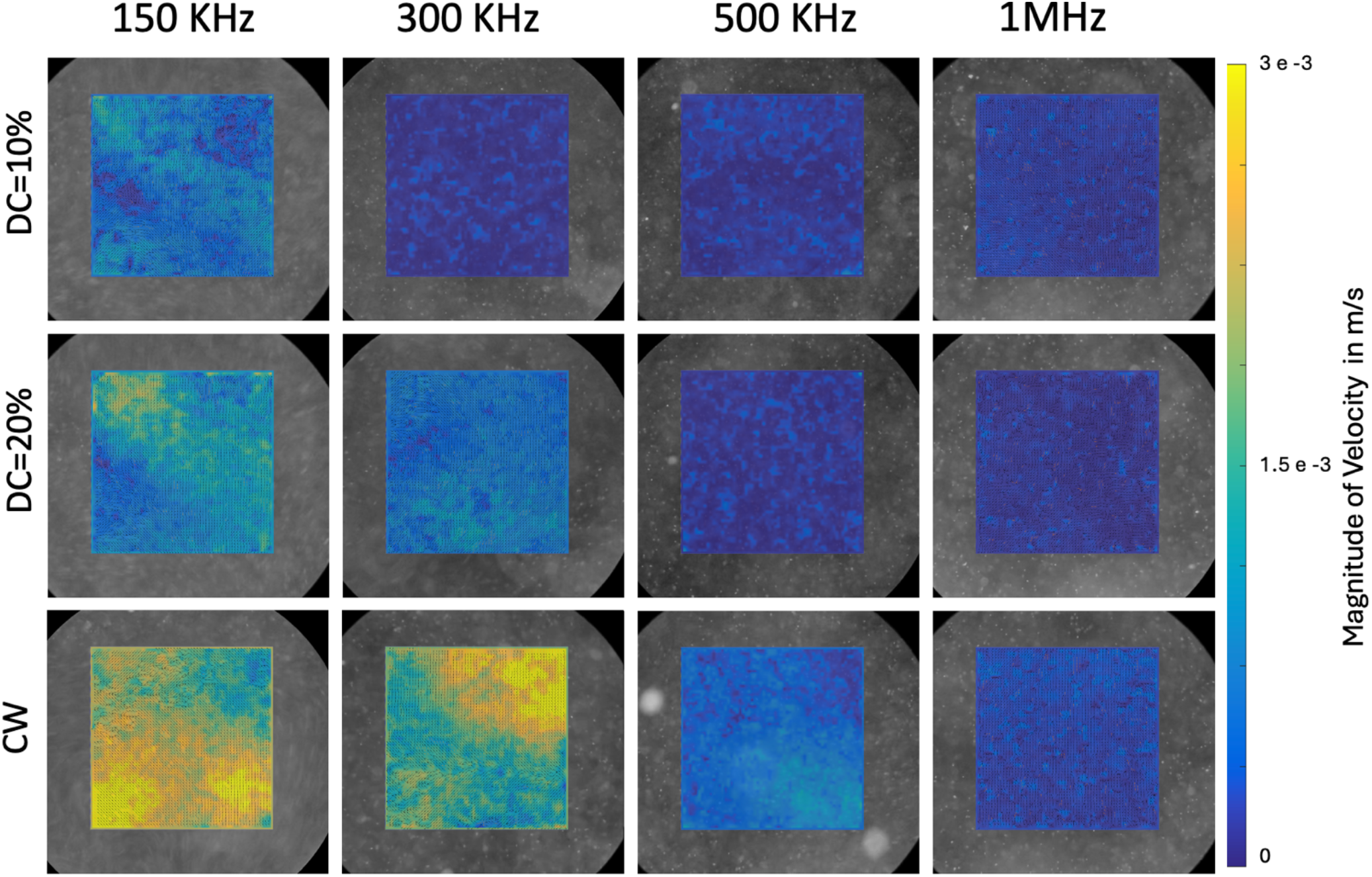
Representative velocity magnitude maps recorded at four acoustic excitation frequencies (150 kHz, 300 kHz, 500 kHz, and 1 MHz) and three duty cycles (10%, 20%, and CW operation). Velocity fields are superimposed on the bright-field image of the observation chamber. Increasing duty cycle consistently amplified streaming velocities, with the highest values observed under CW excitation. The measurements are done at driving pressure of 300 kPa. The velocity scale bar (right) reflects absolute magnitude in m/s.

At 150 kHz, substantial streaming is evident even at 10% duty cycle, with a broad region of elevated velocities across the interrogation area. Increasing duty cycle to 20% and CW further enhances the overall velocity level, and majority of the field of view reaches the upper end of the color scale (3×10^-3^ m/s), indicating strong flow throughout the chamber region sampled by PIV. At 300 kHz, velocities are lower at 10% duty cycle, but increase visibly at 20% and CW, where a large fraction of the interrogation window shows intermediate-to-high velocities.

In contrast, at 500 kHz the velocity maps remain predominantly less than 3×10^-4^ m/s (dark blue) at 10% and 20% duty cycle, indicating very low streaming, with only modest increases under CW excitation. At 1 MHz, the maps are uniformly less than 3×10^-4^ m/s (dark blue) for all duty cycles, consistent with very weak streaming and minimal bulk motion of tracer particles within the measurement region. Overall, Figure 2 illustrates that both frequency and duty cycle strongly influence the magnitude and spatial extent of microstreaming, with low-frequency, high-duty-cycle conditions producing the largest and most widespread velocities. At 500 kHz, the velocity fields are dominated by values near the lower end of the scale at 10% and 20% duty cycles, consistent with weak streaming. CW excitation produces some increase in flow magnitude, but the streaming remains modest and confined compared with the lower frequencies. At 1 MHz, nearly the entire interrogation window exhibits velocity magnitudes near the minimum of the color scale under all duty cycles. The velocity map indicates very weak microstreaming and minimal bulk displacement of tracer particles.

Across all frequencies, increasing duty cycle consistently amplifies the streaming response. Since acoustic pressure was held constant within each frequency condition, the observed differences reflect true frequency and duty-cycle dependent microbubble behavior rather than variations in acoustic input power. Overall, Figure 2 demonstrates that microstreaming is strongest at 150 kHz and 300 kHz frequencies, and becomes progressively weaker as the frequency increases.

### Shear-rate fields

Shear rate represents the spatial gradient of the streaming velocity field and quantifies how rapidly adjacent fluid layers deform relative to one another. In microbubble-mediated acoustic flows, shear arises from the strong velocity differentials created by steady microstreaming vortices. Because shear is the mechanical force most directly experienced by biological species such as cells, extracellular matrices, and clot structures, it is widely considered a key mediator of microbubble-enhanced biological effects including membrane permeabilization, increased mass transport, and enhanced thrombolysis. Therefore, shear-rate maps provide a more mechanistic assessment of the bubblefluid interaction than velocity magnitude alone. Figure 3 shows representative shear-rate fields computed from the PIV-derived velocity gradients for all four frequencies and three duty cycles at 300 kPa. As with the streaming velocity fields, the shear-rate distributions show a strong dependence on driving frequency. At 150 kHz, particularly under CW excitation, the shear-rate field displays markedly elevated spatial variations, with large regions having shear greater than 150 s^-1^. Under 150 kHz CW excitation, the shear-rate field shows pronounced excursions toward both the positive and negative extrema of the scale, reflecting strong bidirectional deformation within the microstreaming flow. These large-amplitude gradients are consistent with the intense streaming velocities observed in the corresponding velocity maps. At 300 kHz, shear rate shows a homogenous distribution close to zero at 10% and 20% duty cycles, but becomes visibly structured for CW sonication. In contrast, at 500 kHz and 1 MHz, the shear maps are dominated by values clustered near zero, with only modest fluctuations. Across all duty cycles, these higher-frequency conditions produce relatively uniform fields with limited positive or negative shear excursions, indicating comparatively weak deformation of the fluid. Across the entire dataset, the shear-rate maps reveal the same monotonic trend observed in the velocity fields: lower frequencies and higher duty cycles yield stronger, more spatially extensive shear, whereas higher frequencies generate weaker, limited shear. Because shear is the principal mechanical output of microstreaming, these observations reinforce the conclusion that steady, low-frequency excitation optimally drives microbubblefluid coupling.

**Figure 3.**
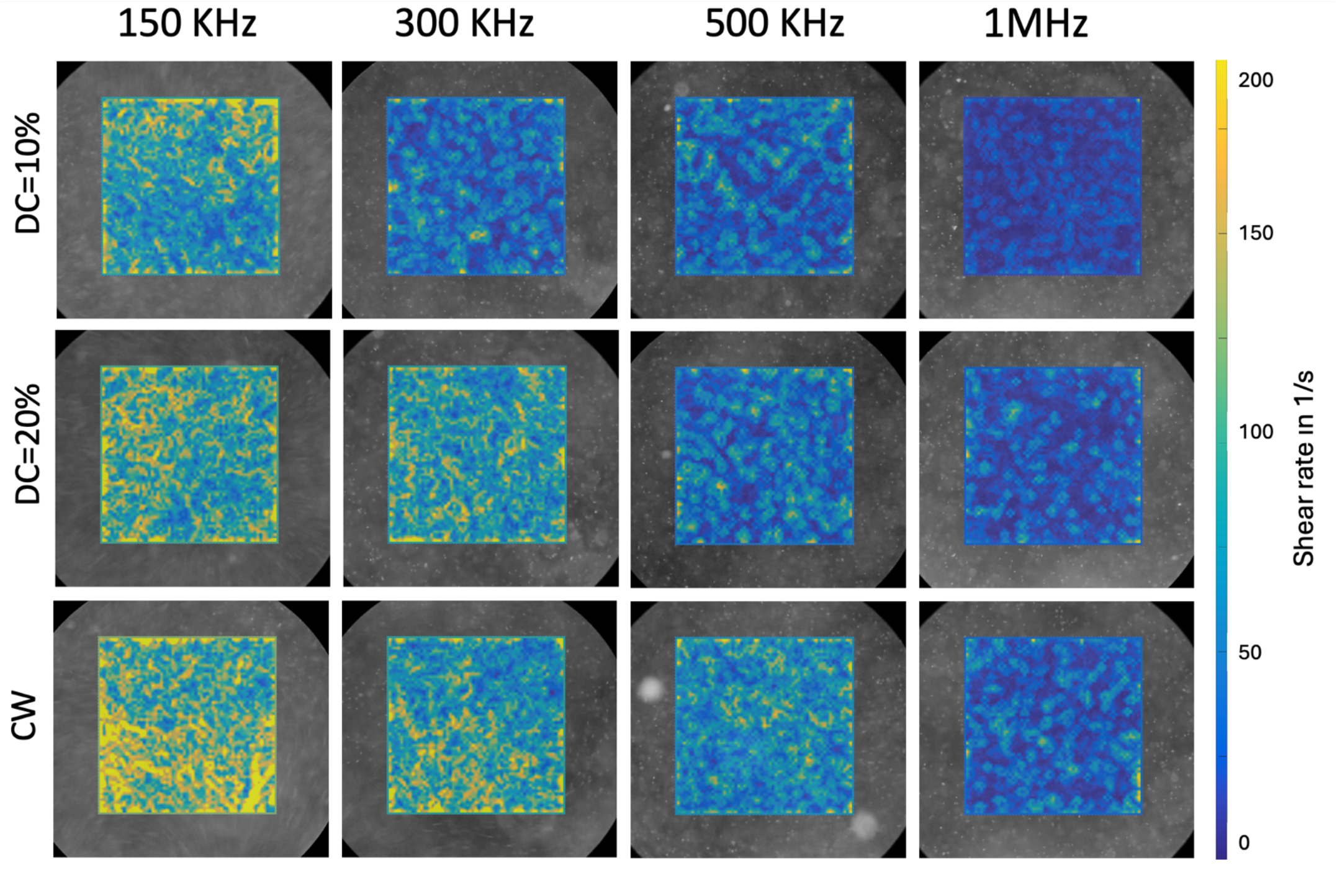
Shear rate fields derived from the PIV vector maps acquired at 150 kHz, 300 kHz, 500 kHz, and 1 MHz under 10%, 20%, and CW excitation. Higher duty cycles produced more distinct Shear rate structures, particularly at 150 kHz and 300 kHz, consistent with stronger microstreaming circulation. At higher excitation frequencies (500 kHz1 MHz), Shear rate patterns remained weaker and spatially homogenous. Shear rate scale is shown at right (in s^−1^).

### Quantitative streaming velocity analysis

While the velocity-magnitude maps in Figure 2 qualitatively illustrate the strong dependence of streaming on frequency and duty cycle, a quantitative analysis is required to compare streaming performance across the full acoustic parameter space. The PIV-derived velocity fields were therefore averaged spatially to obtain mean streaming velocities for each combination of frequency, pressure, and duty cycle. These measurements show how acoustic input conditions modulate the overall strength of microstreaming beyond the spatial patterns seen in the maps.

To compare streaming strength across acoustic conditions, we extracted the mean velocity magnitude from the PIV fields for each combination of frequency, pressure, and duty cycle, averaging the velocity over 100 consecutive frames to obtain a stable measure of steady streaming. Figure 4 shows the resulting mean velocity values across all tested acoustic parameters.

**Figure 4.**
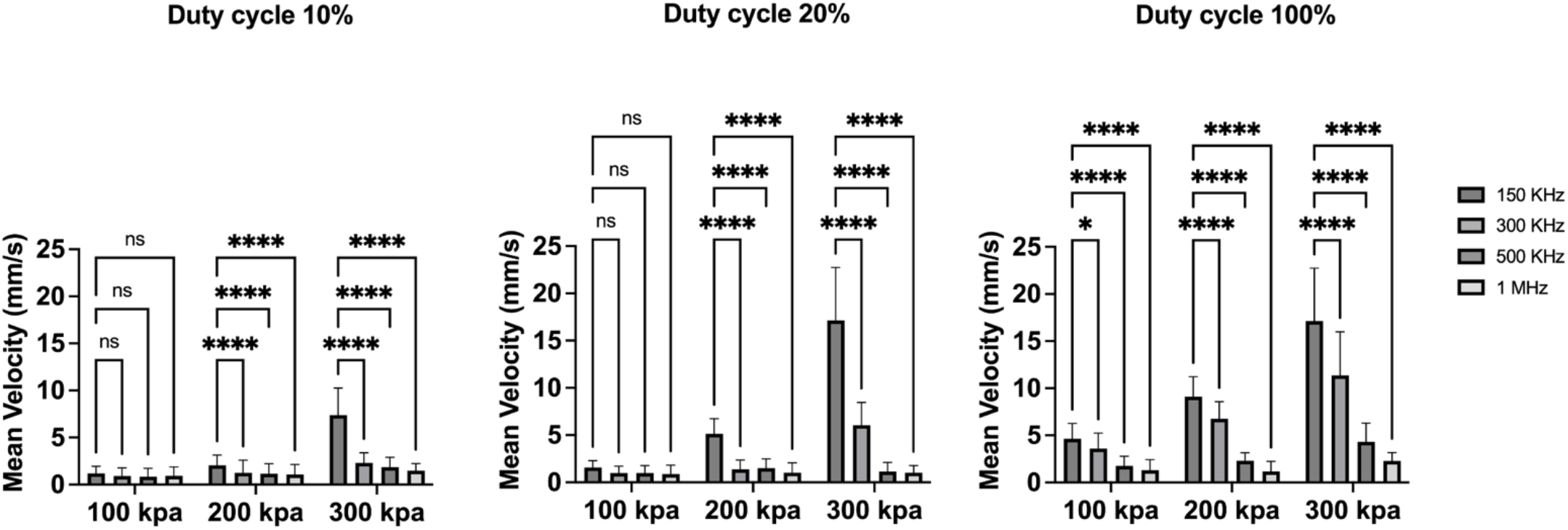
Mean streaming velocity quantified at 100, 200, and 300 kPa. Velocity increased significantly at lower frequencies and at higher pressures, with CW excitation producing the largest rise. Data shown as mean ± SD; multiple-comparison statistics indicated above bars.

Across all duty cycles, mean streaming velocity showed a pronounced dependence on ultrasound frequency. At 150 kHz, mean velocities were consistently the highest, reaching approximately 4e-3 m/s at 100 kPa, 8e-3 m/s at 200 kPa, and 17e-3 m/s at 300 kPa for continuous wave excitation. These values were an order of magnitude larger than those at all higher frequencies. At 300 kHz, mean velocities remained measurable but lower, ranging from ∼3e-3 m/s at 100 kPa to ∼10e-3 m/s at 300 kPa depending on duty cycle. In contrast, 500 kHz and 1 MHz produced only weak streaming, with values generally ≤5e-3 m/s across all pressures and duty cycles. At all duty cycles, the frequency ordering remained similar in that 150 kHz > 300 kHz > 500 kHz > 1 MHz.

Duty cycle strongly modulated these trends. At 10% duty cycle, velocities at 150 kHz increased from ∼1e-3 m/s at 100 kPa to ∼7 e-3 m/s at 200300 kPa). At 20% duty cycle, velocities increased sharply from **∼**3e-3 m/s at 100 kPa to ∼17 e-3 m/s at 300 kPa. Continuous-wave excitation produced the strongest streaming overall and at 150 kHz, mean velocities reached ∼7e-3 m/s at 100 kPa, and ∼17e-3 at 200 kPa**-**300 kPa. The rest of frequencies showed similar trend of increase in intensity by increase in duty cycle, albeit with a smaller range. Acoustic pressure further influenced streaming magnitude. Raising pressure from 100 kPa to 200 kPa and 300 kPa significantly increased mean velocity for every frequency and duty cycle. However, the magnitude of these increases was strongly frequency-dependent. At 150 kHz, pressure doubled or tripled the mean velocity, whereas at 1 MHz, pressure increases produced only minimal changes, consistent with weak bubble fluid interaction at that frequency.

Overall, these quantitative results reinforce the trends observed in the velocity field maps in that low-frequency ultrasound produces substantially stronger microstreaming, and this effect is further enhanced by higher duty cycles and greater acoustic pressures.

## Discussion

Although classical resonance models predict maximal microbubble response near the low MHz range for commercially available contrast-agent of 1-5 micron, our particle image velocimetry measurements reveal pronounced microstreaming velocities and sustained flow structures at sub-MHz frequencies, with the strongest response observed in the 150-300 kHz range. This behavior does not follow the classical expectation that proximity to the nominal resonance frequency necessarily maximizes functional microbubble effects. Our observations are consistent with prior reports demonstrating that low-frequency sonication can drive large-amplitude, stable oscillations in shelled microbubbles even when operating well below resonance. Notably, in one report enhanced microbubble contrast agent oscillation following 250 kHz sonication was reported to be over 16 times higher compared with 1 MHz under similar pressure conditions. ^13^These findings indicate that oscillation amplitude can be substantially larger at low frequency, despite being far from resonance. At lower frequencies, the longer acoustic period permits greater volumetric oscillation per cycle and reduced cycle-to-cycle damping, enabling more effective momentum transfer from the oscillating bubble to the surrounding fluid. In addition, the viscous boundary layer thickness which is the driving force of Rayleigh streaming is known to increase with decreasing frequency, allowing oscillatory motion to penetrate farther into the fluid and promoting stronger steady streaming. ^10^

The strong frequency dependence of microstreaming observed in this study has direct implications for microbubble-mediated biomedical applications in which fluid transport and shear are the primary therapeutic mechanisms. In sonothrombolysis, enhanced clot breakdown has been consistently reported at low ultrasonic frequencies, including in our prior work, where improved lytic efficacy was observed in the same sub-MHz range that here produces the strongest microstreaming.^25^ The present PIV measurements provide a mechanistic explanation for these outcomes: stronger low-frequency streaming generates sustained convective transport and elevated shear near the clot surface, facilitating the mechanical disruption of fibrin networks without requiring inertial cavitation. In this context, frequency-dependent microstreaming offers a more direct predictor of lysis efficiency than cavitation amplitude alone.

Similar trends are evident in blood brain barrier opening studies using focused ultrasound systems operating in the 220-250 kHz range.^17^ Several preclinical and clinical studies have demonstrated reliable BBB disruption using focused ultrasound in the 220250 kHz range in combination with microbubbles, including work by McDannold *et al*. employing 220 kHz. ^17,26^ The present findings suggest that enhanced BBB permeability at these frequencies may, in part, be mediated by increased microstreaming-induced shear and convective transport at the vessel wall, rather than by proximity to microbubble resonance cavitation events.

Taken together, these observations argue that low-frequency ultrasound is intrinsically more effective at harnessing microbubble-driven microstreaming for functional applications, including clot lysis and barrier disruption. While pressure and duty cycle can scale the magnitude of the response, they do not compensate for inefficient bubblefluid coupling at higher frequencies. Thus, when therapeutic efficacy relies on streaming-driven transport and mechanical interaction, selecting an appropriate low-frequency regime is likely more critical than operating near the nominal resonance frequency of the microbubble.

## Conclusion

In this study, we directly measure the interaction between acoustically driven microbubbles and the surrounding fluid using particle image velocimetry, rather than inferring activity from bubble oscillation amplitude or acoustic emissions. This approach allows direct quantification of microstreaming velocity and shear, which are the primary mechanical mechanisms underlying many functional microbubble therapies. This work demonstrates that microbubble-driven microstreaming is highly sensitive to ultrasound frequency, with low-frequency excitation (150-300 kHz) producing substantially stronger and more spatially extensive streaming than higher-frequency ultrasound. The strong agreement between our PIV-derived velocity and shear fields and previously reported low-frequency microbubble oscillation behavior supports the conclusion that sub-MHz excitation can generate substantial microstreaming through stable cavitation, without reliance on resonance tuning or inertial cavitation.

Increasing duty cycle and acoustic pressure amplify streaming magnitude but do not alter the fundamental frequency dependence. Particle image velocimetry reveals that the resulting shear fields which is critical to many microbubble-mediated therapeutic mechanisms follow the same monotonic trend, with 150 kHz generating the highest shear levels across all tested pressures and duty cycles.

These results establish a quantitative framework for selecting acoustic parameters that optimally harness microbubble-induced streaming. By identifying low-frequency, stable-cavitation regimes that maximize velocity and shear generation, this study provides guidance for improving the efficacy of microbubble-enabled applications such as drug delivery, clot disruption, and barrier permeabilization. The findings underscore the importance of direct flow-field measurement in evaluating microbubble functionality and highlight the potential for PIV-informed optimization of ultrasound-based biomedical therapies.

